# Thermal Conductivity of Artificial Materials Engineered from Plant and Bacterial Cells

**DOI:** 10.64898/2026.05.04.722776

**Authors:** Masaki Odahara, Yoko Horii, Tashi Xu, Kayo Terada, Kazuho Daicho, Junichiro Shiomi, Keiji Numata

## Abstract

Bio-based materials are known for their excellent biodegradability and, in some cases, their potential to fix carbon dioxide. Owing to these properties, they are increasingly being utilized as environmentally friendly alternatives across various applications. In this study, we focused on using living cells themselves as material components, aiming to evaluate their potential as substitutes for conventional plastic-based thermal insulators. We selected two types of cells, photosynthetic purple non-sulfur bacterium *Rhodovulum sulfidophilum* and tobacco BY-2 plant suspension cells. After optimizing solidification conditions through the addition of pectin and cellulose nanofibers, we measured the thermal conductivity of the solidified cells under atmospheric pressure. The results showed that *R. sulfidophilum* exhibited 0.0553 W/m·K, while BY-2 exhibited a thermal conductivity of 0.043 W/m·K. Both values indicate relatively low thermal conductivity compared to existing bio-based materials, suggesting high insulation performance. Among the solidified cells, the solidified BY-2 cells showed minimal variation in thermal insulation performance under pressure changes, and had a low thermal emissivity as revealed by FT-IR analysis. Based on these findings, we propose that cell-derived materials can serve as potentially biodegradable bio-based thermal insulation materials.

## Introduction

Biological materials are utilized in various aspects of our daily lives. Among them, plant-derived materials such as wood, fibers, and rubber are environmentally friendly. In particular, wood, which exhibits excellent carbon dioxide fixation capacity and biodegradability, has been attracting renewed attention as an eco-friendly construction material [1]. On the other hand, thermal insulation materials used inside buildings require molding, and thus plastic-based fillers such as polystyrene are widely used. Therefore, the development of environmentally friendly alternatives is strongly desired.

We focused on photosynthetic bacterial and plant suspension cells as environmentally friendly biological materials that could potentially serve as candidates for thermal insulation materials. One of them is the marine purple non-sulfur bacterium *Rhodovulum sulfidophilum*, a Gram-negative species [2]. The cells are spherical, approximately 1–3 μm in diameter, and grow in a dispersed form under standard culture conditions. *R. sulfidophilum* is a photosynthetic bacterium that can grow under a wide range of conditions, from photoautotrophic to heterotrophic [3], and possesses the ability to fix both carbon dioxide and nitrogen [4]. With this characteristics, *R. sulfidophilum* can be used as biomass for sustainable fertilizer for plant production [5].

The other cell type is the tobacco (*Nicotiana tabacum*)-derived cultured cell line BY-2 [6]. BY-2 cells are elongated and cylindrical, measuring 30–100 μm in length, surrounded by a cell wall [7], and typically form bead-like chains consisting of about 2– 8 cells. BY-2 exhibits a faint yellow coloration because of lack of photosynthetic ability. BY-2 can be cultured in liquid medium, proliferates rapidly, and has been used not only as a model system for fundamental studies of plant cell biology but also as a platform for protein production [8, 9].

In this study, we evaluated the potential of *R. sulfidophilum* and BY-2 cells themselves as structural materials (Figure 1). After solidifying the cells with the aid of additive agents, we conducted structural and optical assessments, which revealed distinct characteristics for each type of solidified cell. Based on these properties, we propose that solidified cells can serve as novel biodegradable materials for thermal insulation.

**Figure 1.**
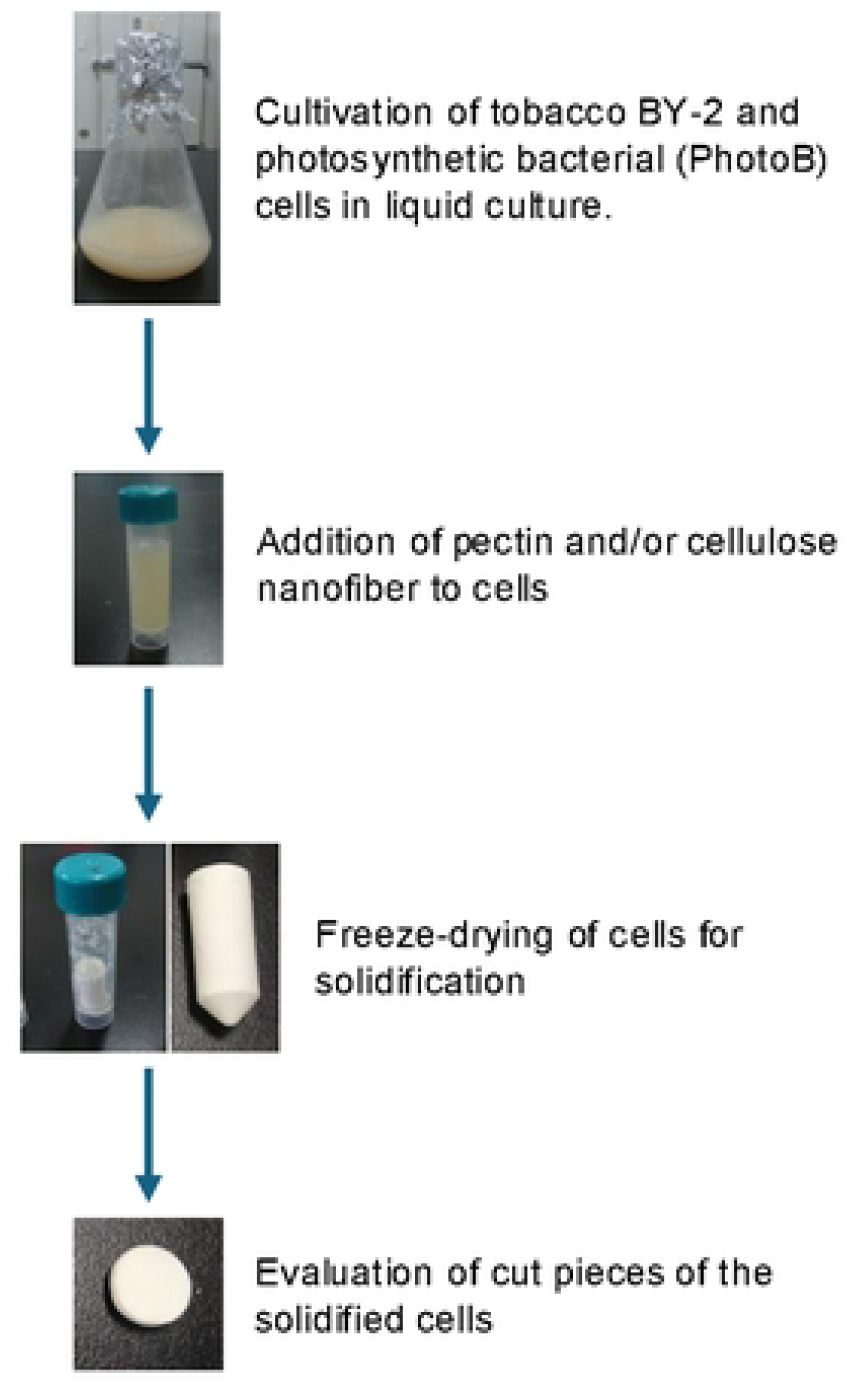
Sample preparation scheme of the study.

## Materials and Methods

### Organisms and culture conditions

*Nicotiana tabacum* BY-2 cell was cultivated by rotary shaker (130 rpm) in a modified Linsmaier and Skoog (mLS) liquid media, pH 5.8 supplemented with 0.2 mg L^−1^ 2,4-dichrorophenoxyacetic acid (2,4-D) and 30 g L^−1^ sucrose in the dark at 27°C. *Rhodovulum sulfidophilum* (ATCC 35886) was cultivated in marine broth (Merck, Germany) at 26°C under 80 W m^-2^ irradiation, according to Morey-Yagi, Hanh (10).

### Solidification of BY-2 and PhotoB cells

For BY-2 cells, after pelleting 7-day-culture (3-5×10^6^ cells/mL) by allowing the tubes to stand for 10 minutes, the culture medium was removed. The cells were washed three times with pure water, then 3 g of cells were mixed with pectin (FUJIFILM Wako, Japan) and cellulose nanofiber. After centrifugation at 2,000 × g for 1 minute to remove air, the cells were solidified using a lyophilizer (FDU-2100, EYELA, Japan). For PhotoB, cells from 6-day culture with OD_660_=~2.5 were harvested by centrifugation at 14,000 × g for 10 minutes and washed once with pure water. After a second centrifugation at 14,000 × g for 30 minutes, 1 g of the cells were solidified using the same lyophilizer (FDU-2100, EYELA, Japan).

### Scanning electron microscopy

Solidified cell materials were sputtered with gold for 1 min using a JEOL Smart Coater (JOEL, Japan) and then observed by JCM-6000 (JEOL, Japan).

### Measurement of thermal conductivity

Thermal conductivity of materials was measured using a steady-state setup shown in Figure 4a following the steady-state method reported by Obori, Suh (11). In solidified cell steady-state measurements, the thermal conductivity is obtained directly from Fourier’s law, and thus no specific sensitivity is defined. The measurement accuracy is mainly governed by the uncertainty in the temperature difference across the sample, and a sufficiently large temperature gradient is required. In this work, a temperature difference exceeding 10 K was maintained to ensure reliable measurements. Cell-based solidified materials were cut into ~6.5 mm thickness with 78.5 mm^2^ cross section for solidified BY-2 and 2.1-2.3 mm thickness with 78.5 mm^2^ cross section for solidified PhotoB by a razor. A sample was set between the copper blocks, the steady-state temperature difference was imposed between the upper and lower side of the sample, and the temperatures and heat flux were measured with the sensors to extract thermal conductivity based on Fourier’s law. The system was cased in a vacuum chamber to vary the pressure, as the vacuum level in this range can directly affect heat transfer in porous materials. For humidity, both low- and high-vacuum results should not be affected by the variation of humidity. Measurements at atm were conducted in an air-conditioned room, within the same season. During these measurements, the sample was still placed inside the chamber to minimize potential effects from ambient humidity. Calibration was performed to adjust the sensor’s output current to match the ambient temperature. Polymeric form with styrene (5 mm thickness) and glass wool insulation (3 mm thickness) (Daytone, Japan) were used as references.

### FT-IR

The Fourier-transform infrared (FT-IR) spectra of each sample were recorded on an IRPrestige-21 FT-IR spectrophotometer (Shimadzu Corporation, Kyoto, Japan) with a MIRacle A single-reflection attenuated total reflectance (ATR) unit using a Ge prism. The spectra from 700 to 4000 cm^− 1^ were accumulated at 4 cm ^−1^ resolution using 128 scans. ATR correction was carried out at 1600 cm^-1^ and 1650 cm^-1^ for BY-2 and PhotoB cells, respectively.

## Results and Discussion

### Solidification of photosynthetic bacterial and plant cells

For solidification of the cells, we tested freeze-drying. The cells were harvested and washed with water to remove residual culture medium, followed by freeze-drying. However, the freeze-dried BY-2 cells exhibited a fluffy texture that hindered accurate measurement of thermal conductivity (Figure 2a). To improve the compactness of the samples, we added pectin and/or cellulose nanofiber (CNF) as supportive agents, aiming to promote cell–cell adhesion and act as a filler, respectively. The addition of pectin alone hardened the BY-2 cells, but the resulting materials remained fragile even with 5% pectin (w/w relative to cell weight) (Figure 2a). We next applied CNF at 25% and 50% (w/w), and found that 25% CNF combined with 1% pectin sufficiently increased the hardness of the solidified cells, making them suitable for subsequent analyses (Figure 2a). Since increasing CNF to 50% did not cause further qualitative changes, we used 1% pectin and 25% CNF as the optimal additives for solidification in the following experiments. Note that the solidified BY-2 cells exhibited a white color even without CNF, likely because their undifferentiated plastids lack photosynthetic ability and therefore do not accumulate pigments [12].

**Figure 2.**
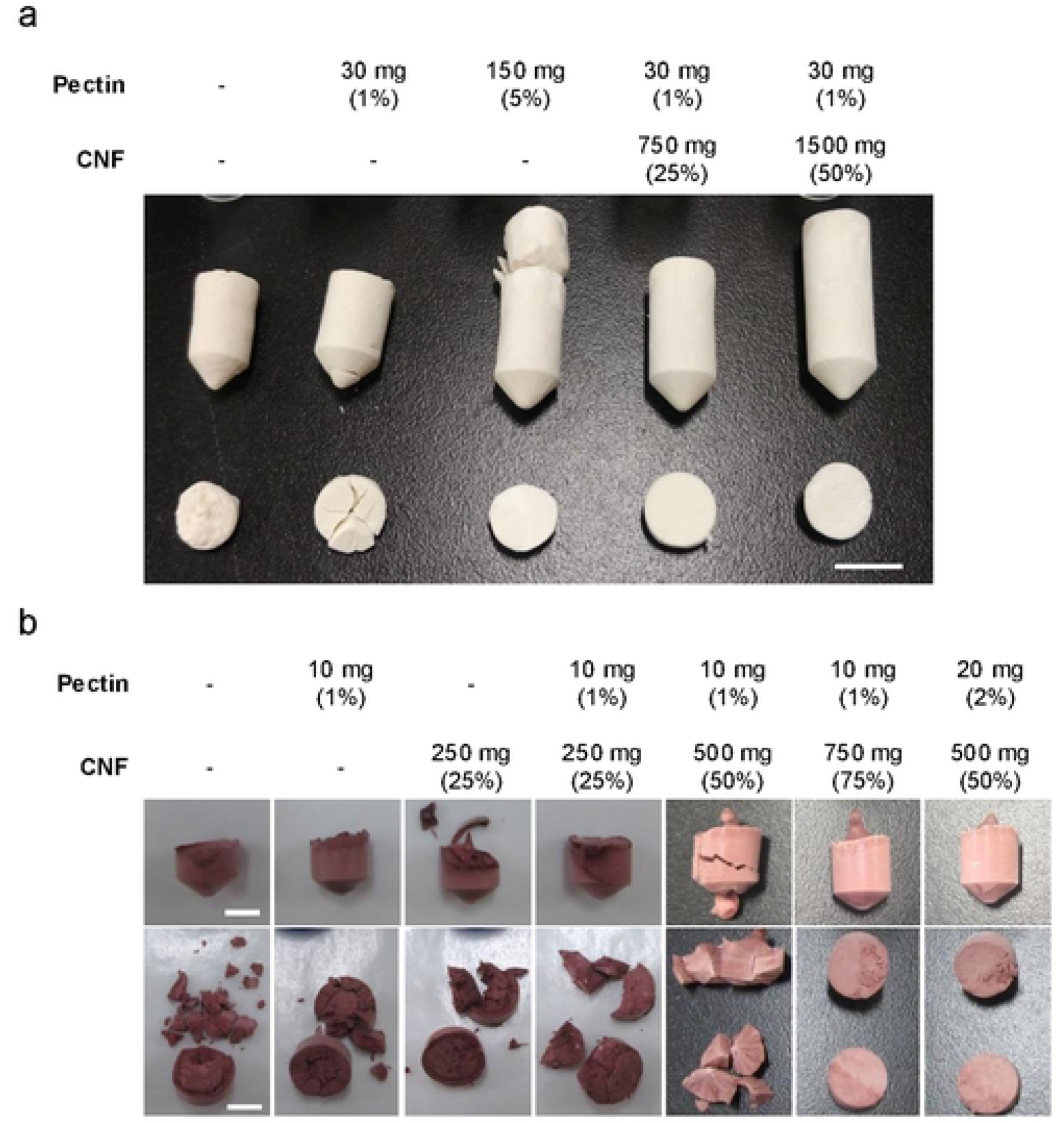
Optimization of additive agents for solidification of plant and photosynthetic bacterial cells. For solidification, pectin and/or cellulose nanofiber (CNF) were added to 3 g dry weight (DW) of BY-2 cells (a) or 1 g of photosynthetic bacterial cells (b) in the amounts indicated above the images. Cut sections of the solidified cells are shown in the lower parts of the images. Bars, 10 mm in (a) and 5 mm in (b).

Similarly to the BY-2 cells, we optimized the solidification conditions for PhotoB cells. As with BY-2 cells, the freeze-dried PhotoB cells were fragile and unsuitable for thermal conductivity analysis (Figure 2b). The addition of either or both 1% pectin and 25% CNF did not sufficiently solidify the PhotoB cells (Figure 2b), in contrast to the case of BY-2 cells. This difference may be attributed to the clustering of BY-2 cells, which likely facilitates solidification. Although increasing the CNF content to 50% did not achieve sufficient solidification of PhotoB cells, the addition of 75% CNF did. Alternatively, PhotoB cells were also successfully solidified by combining 2% pectin with 50% CNF. Each of the solidified cell maintained their integrity for more than 17 months under the lab condition (Figure S1), suggesting that they are not fragile. The density of the solidified cells prepared with 1–2% pectin and 50% CNF was 0.037 g/cm^3^ for BY-2 and 0.099 g/cm^3^ for PhotoB. These optimizations enabled further analyses of the solidified cells and revealed differences in the physical nature of the two types of freeze-dried cells.

### Structural assessment of the solidified cells

To investigate the structural properties of the solidified cells, particularly pore characteristics and the integrity of the original cell structures, we conducted scanning electron microscopy (SEM) analysis. SEM revealed that solidified BY-2 cells without additives exhibited rough surfaces with large pores (Figure 3a). The pore sizes roughly corresponded to the typical BY-2 cell diameter (30–100 µm)[13], suggesting that the pores primarily originated from collapsed or cavitated cells. The addition of pectin slightly smoothed the surface, although pores remained visible (Figure 3a). In contrast, the addition of CNF produced a much smoother surface with fewer pores, indicating that CNF likely filled the pore spaces (Figure 3a).

**Figure 3.**
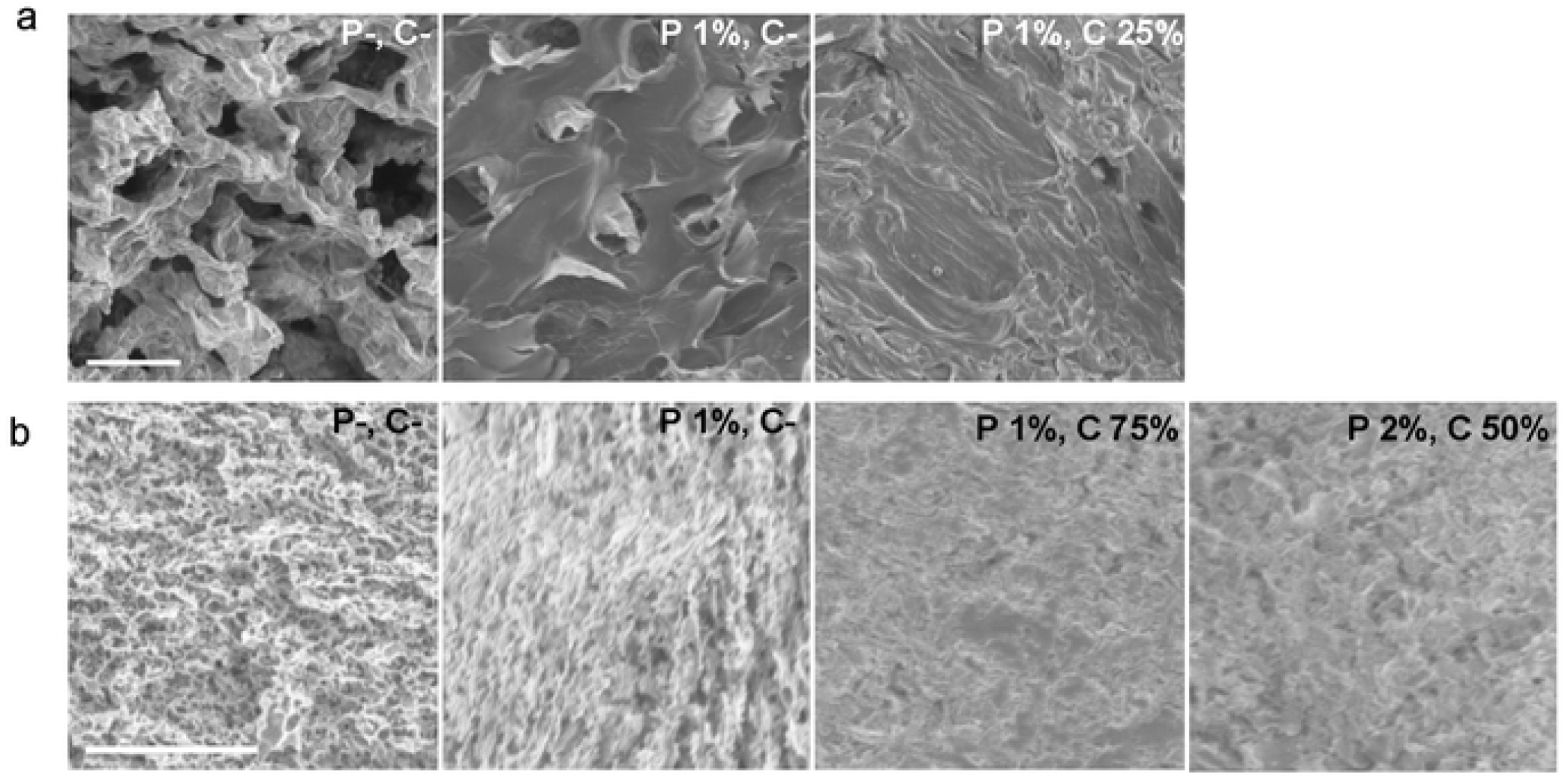
Surface of the solidified plant and photobacterial cells observed by SEM. a. SEM images of the BY-2 cells solidified with pectin only or with pectin and CNF. b.SEM images of the PhotoB cells solidified with only or with pectin and CNF. P and C denote pectin and cellulose nano fiber, respectively. Bars, 50 μm

**Figure 4.**
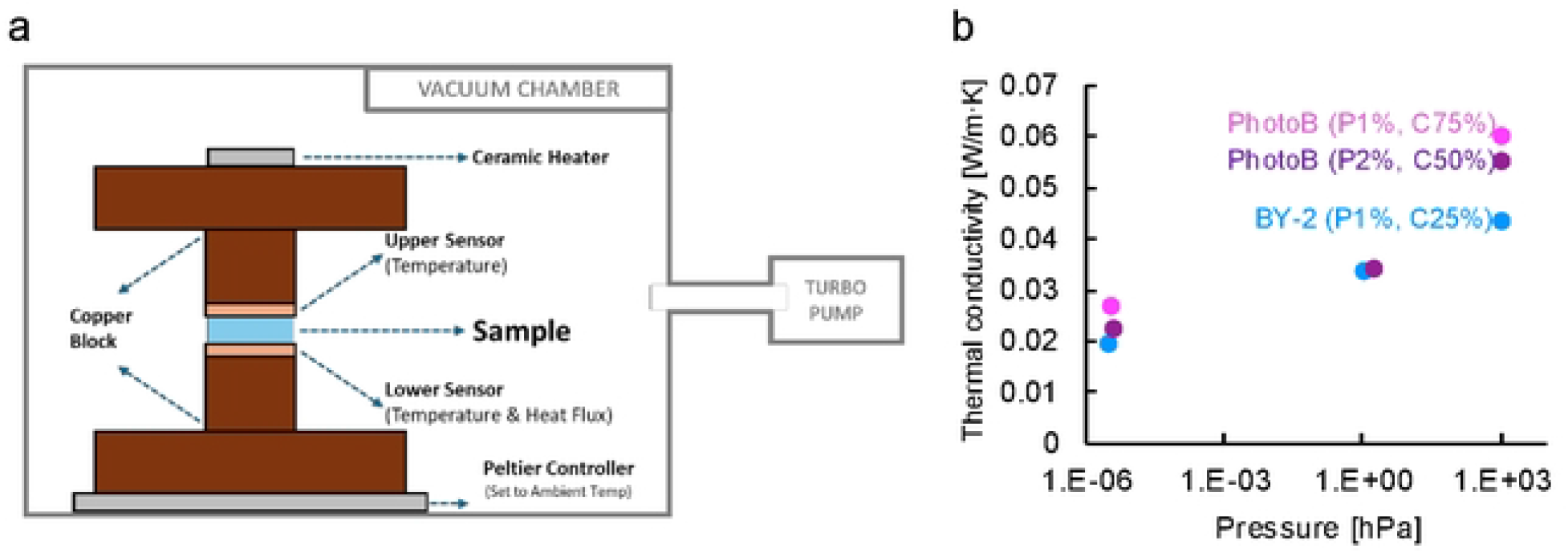
Thermal conductivity of the solidified cells. a. Scheme for measurement of thermal conductivity. Samples are placed between the thermocouple, and thermal conductivity is measured under the control of pressure. b. Thermal conductivity of the three types of the solidified cells at different pressures.

Compared with solidified BY-2 cells, solidified PhotoB cells without additives exhibited smaller pores and lower surface roughness (Figure 3b), consistent with their smaller cell size (1–3 µm). The addition of pectin alone caused no noticeable change in the surface features, and small pores remained visible (Figure 3b). Under sufficiently solidified conditions (+1% pectin and 75% CNF, or +2% pectin and 50% CNF), the surface became smoother and pores were scarcely observed, likely due to pore filling by CNF as observed in the BY-2 samples (Figure 3b). Overall, SEM analysis showed that cell morphology was largely lost after processing and highlighted the differences in pore size between solidified BY-2 and PhotoB cells, reflecting their intrinsic cell sizes, and proposed that most pores were likely filled with CNF.

### Thermal conductivity of the solidified cells

The porous characteristics of the solidified cells revealed by SEM suggest high thermal insulation properties. Thermal conductivity was evaluated for solidified BY-2 and PhotoB cells under different pressure conditions as shown in the Figure 4a. The solidified BY-2 cells (+1% pectin, 25% CNF) exhibited a thermal conductivity of 0.0426 W/m·K at atmospheric pressure (1000 hPa), whereas the solidified PhotoB cells showed 0.0603 W/m·K for the +1% pectin, 75% CNF sample and 0.0553 W/m·K for the +2% pectin, 50% CNF sample (Figure 4b). These results indicate that the solidified BY-2 cells had lower thermal conductivity than the solidified PhotoB cells at atmospheric pressure. All measured values were lower than those of wood, whose minimum thermal conductivity is approximately 0.079 W/m·K [14]. Even compared with primary cell walls of onion tissue, which exhibits a minimum of 0.29 W/m·K across dry to wet conditions [15], both types of solidified cells displayed substantially lower thermal conductivity.

Under vacuum conditions, all solidified samples showed decreased thermal conductivity at around 1 hPa compared with 1000 hPa, and even lower values at 3–4 × 10^−6^ hPa (Figure 4b). Notably, the solidified BY-2 cells consistently exhibited the lowest thermal conductivity across all pressure ranges and the smallest variation among pressures. These characteristics suggest that the solidified cells contain open pores that form air pathways within the material, and that the BY-2 cells themselves have lower intrinsic thermal conductivity than the PhotoB cells. In this regard, the result that the thermal conductivity of BY-2 cells measured under as similar conditions as possible was lower than that of PhotoB (Table S1) suggests that solidified BY-2 cells possess superior thermal insulation properties. The cellulose composition of the BY-2 cell wall may contribute to this lower thermal conductivity. On the other hand, the increase in thermal conductivity toward approximately 0.07 W/m·K at atmospheric pressure with increasing CNF content in solidified PhotoB suggests that CNF has a higher thermal conductivity than PhotoB cells, which is consistent with the reported thermal conductivity of CNF (0.06–0.07 W/m·K) [16]. Pectin, the other additive, is present only in a small amount, and thus its effect on thermal conductivity is considered to be very limited.

To further examine the pressure-dependent dynamics of thermal conductivity in the solidified BY-2 cells, we compared their behavior with those of reference porous materials. Thermal conductivity was measured for a polymeric foam (representing a closed-porous material) and high-density glass wool (representing open-porous materials) under various pressures. All materials showed little change in thermal conductivity below approximately 0.01 hPa, followed by an increase with rising pressure above this threshold. However, the patterns of increase differed between closed- and open-porous materials (Figure S2). The thermal conductivity of the solidified BY-2 cells roughly followed that of the open-porous materials, suggesting that the solidified BY-2 cells behave as open-porous materials rather than closed ones. Nevertheless, the solidified BY-2 cells are likely more closed than the solidified PhotoB, as indicated by their smaller changes in thermal conductivity across pressures (Figure 4b). This is further supported by their SEM images, which show fewer pores in the solidified BY-2 cells (Figure 3).

Another factor that can contribute to the measured pressure dependence of thermal conductivity is the effect of thermal convection. At the surfaces of the samples that are in contact with the surrounding gas, heat can dissipate to the ambient gas by natural convection, which leads to overestimation of heat flux and thermal conductivity. This effect decreases with pressure and thus can contribute to the pressure dependence of thermal conductivity. The impact of effect scales with the surface to volume ratio of the material, and it should not be significant for the current samples. Nevertheless, it is worth noting that, because the solidified BY-2 sample is thicker than the solidified PhotoB samples, the finding that solidified BY-2 exhibits lower thermal conductivity remains valid even when considering the influence of thermal convection. In addition, the thermal conductivities at atmospheric pressure presented here are the upper limit values including the effect of thermal convection and the actual values can be even smaller.

Among the PhotoB-based solidified cells, the sample containing 1% pectin and 75% CNF exhibited slightly higher thermal conductivity than that with 2% pectin and 50% CNF at certain pressure points (Figure 4b). This difference can be attributed to the relatively high thermal conductivity of CNF (0.06–0.07 W/m·K) [16], suggesting that the PhotoB cells themselves possess lower intrinsic thermal conductivity than CNF. In summary, the solidified cell materials exhibited thermal conductivity values lower than those of natural woods and comparable to typical insulation materials such as polystyrene (~0.035 W/m·K) [17] or polyurethane (0.033-0.034 W/m·K) [18].

### Photochemical property of the solidified cells

Fourier transform infrared (FT-IR) spectroscopy is commonly used to evaluate molecular structure of wood [19]. To assess the molecular structure of the solidified BY-2 and PhotoB cells, FT-IR transmittance spectra were measured. The spectrum of the solidified BY-2 sample (+1% pectin, 25% CNF) closely matched that of the cell-free material (+1% pectin, 25% CNF) (Figure 5a), suggesting that the spectrum of the solidified BY-2 is mainly derived from the cell wall components and that the BY-2 cells themselves exhibit high transmittance, similar to CNF. Because the spectrum of the cell-free material was highly consistent with that of CNF [20], the contribution of pectin to the overall spectrum appears to be minimal. This high optical transmittance of BY-2 cell is explained by losing pigments for photosynthesis [6]. Absence of clear bands around 1513 cm^−1^ 2920-2925 cm^−1^ corresponding to lignin in leaf [21] and woods [19] verifies lack of lignin in the solidified BY-2.

**Figure 5.**
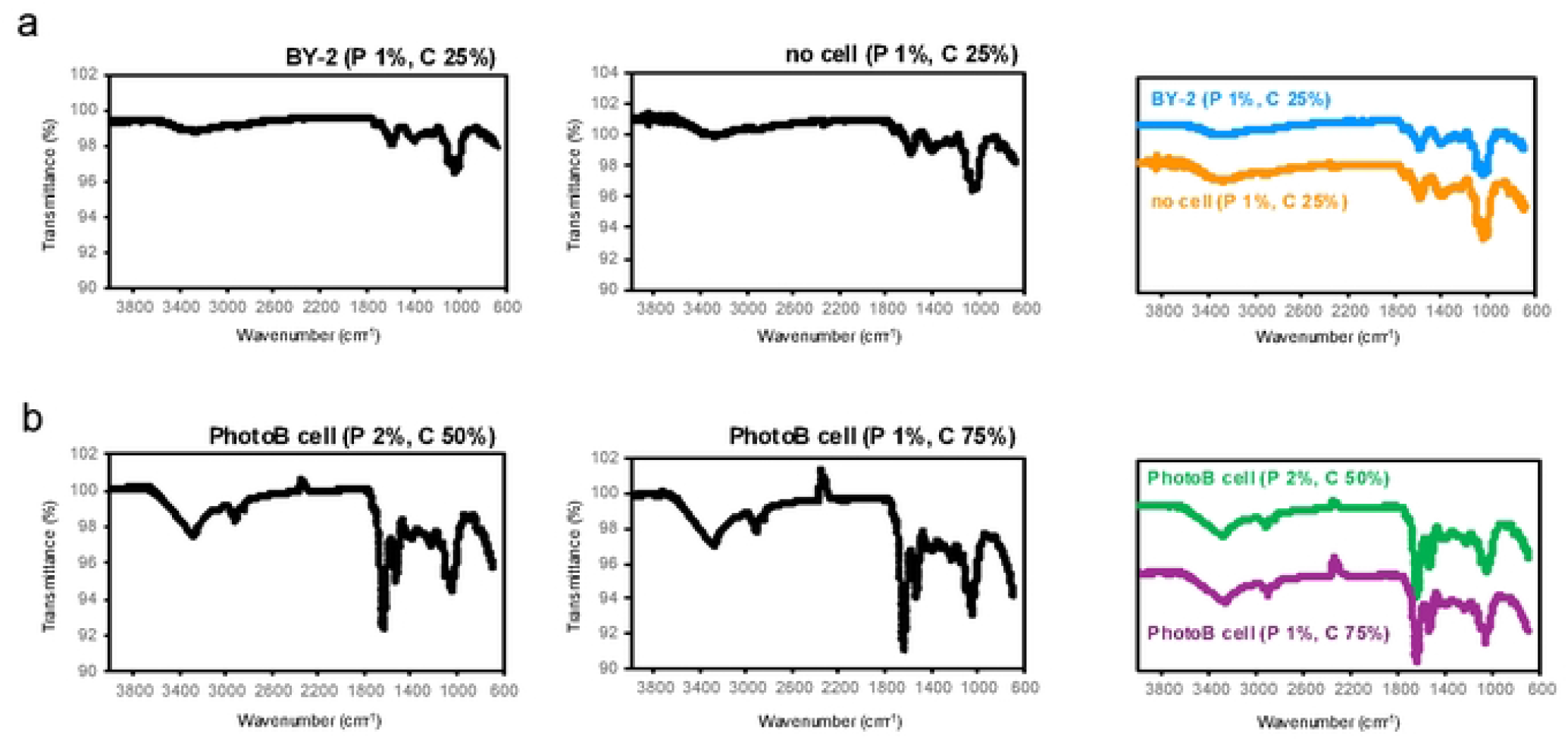
FT-IR measurement of the solidified cells. a. FT-IR spectra of solidified BY-2 cells with 1% pectin and 25% CNF (left), solidified 1% pectin and 25% CNF (middle), and merge of the two spectra (right). b. FT-IR spectra of solidified PhotoB cells with 2% pectin and 50% CNF (left), PhotoB cells with 1% pectin and 75% CNF (middle), and a merge the two spectra (right).

Comparison between the spectra of solidified PhotoB cells and the 1% pectin and 25% CNF sample showed broadly decreased transmittance in the solidified PhotoB spectrum, except for a band at 1000–1150 cm^−1^, which is characteristic of CNF (Figure 5b). The solidified PhotoB cells also exhibited several distinct peaks, particularly at 1650 cm^−1^ and 1550 cm^−1^ (Figure 5b), corresponding to typical bacterial absorption bands [22]. The two types of solidified PhotoB samples showed similar spectral patterns, except for decreased transmittance around 1650 cm^−1^ in the +2% pectin, 50% CNF sample (Figure 5b), which is attributable to the increased amount of PhotoB cells. In summary, the FT-IR spectra of the solidified cells indicated high transmittance in the BY-2 samples and slightly lower transmittance in the PhotoB samples. Given the close relationship between IR absorbance and thermal emissivity, these results are consistent with BY-2 samples exhibiting lower effective thermal emissivity.

## Conclusion

In this study, we successfully solidified and dried two types of cells—photosynthetic bacteria-derived and plant-derived—by adding pectin and CNF as additive, and measured their thermal conductivity. The measured thermal conductivity was lower than that of the additive CNF or woods, and was comparable to that of polystyrene or polyurethane. In particular, for BY-2 cells, the results suggested that the low thermal conductivity of the dried cells was greater than that of PhotoB, and the solidified BY-2 cells exhibited high light transmittance. Furthermore, the BY-2 cell materials show lower thermal conductivity than the bio-based insultation such as composite material containing coir fiber (0.0624 to 0.04628 W/m·K) [23]. Such cell-derived materials, like wood, are both potentially biodegradable and moldable. Given their low thermal emissivity, solidified BY-2 could serve as a “white wood” material with low thermal conductivity, provided that their mechanical strength is sufficiently improved. In contrast, PhotoB materials could contribute to reducing carbon dioxide emissions by environmental carbon fixation through photosynthesis. Overall, these solidified cells represent promising candidates for potentially biodegradable insulating materials. While our solidified cells were revealed unlikely to be fragile under the lab condition, it will be necessary to evaluate the mechanical strength of these solidified cells, as well as their performance under a range of environmental conditions, including humidity, temperature, and light exposure, for practical application as thermal insulation materials. Considering their nature, we propose that these cell-derived materials have potential as sprayed insulation. In addition, given their bio-based nature, further assessments of susceptibility to biological contamination, biodegradability, and long-term stability are required. On the other hand, this study also demonstrated that additives are essential for cell solidification, highlighting a limitation of the current approach. Addressing these challenges, including the optimization of additives, will be critical for advancing the practical use of these materials as thermal insulators.

## Acknowledgements

We thank RIKEN BRC for providing tobacco BY-2 cell.

